# A Practical Framework for Constructing Population-Specific and Alternate-Contig-Aware Genome References: A case study of Vietnam

**DOI:** 10.64898/2026.08.24.746817

**Authors:** Trang T. H. Tran, Vinh C. Duong, Nam N. Nguyen, Tien M. Pham, Quang T. Vu, Mai H. Tran, Tham H. Hoang, Quan Nguyen, Nguyen Thuy Duong, Nam S. Vo

## Abstract

Current studies in human genomics typically rely on the standard genome reference GRCh38 which is known to be biased toward populations of European ancestry and therefore has limitations when applied to other populations. Although various graph-based pangenome references were constructed for several populations to deal with this bias, their usage in practice is currently still limited compared to linear genome references. Here we present a framework for constructing a population-specific genome reference using GRCh38 as backbone with alternate-contig awareness to enhance genomic data analysis in the target population. We demonstrated the advantages of our framework using both public and in-house Vietnamese whole-genome sequencing (WGS) datasets. Genomic variants derived from high-coverage WGS data of the 1000 Vietnamese Genomes Project (VN1K) were imported into our framework to build a Vietnamese-specific Genome Reference (VGR). VGR was then compared to GRCh38 in read alignment and variant calling using high-coverage WGS data of 99 Vietnamese individuals (KHV) from the 1000 Genomes Project (1kGP). Using Omni array genotyping data from 99 KHV samples as an independent benchmark, we found that VGR improved variant-calling precision and reduced false-positive calls compared to GRCh38. Our framework could be easily used for other populations as long as they have a variant database similar to VN1K. Our code is publicly available at github.com/VinGenome/VGR

## 1. Introduction

Whole-genome sequencing (WGS) powered by Next-Generation Sequencing (NGS) technologies enables the comprehensive detection of genetic variants and has been shown to be highly effective in identifying associations between genetic predisposition and diseases(1). In WGS analysis pipelines, the use of a genome reference is essential, as it provides a standardized coordinate framework for systematically documenting and comparing findings across diverse global studies(2). As sequencing technologies advanced, genome references were progressively improved to provide more accurate and comprehensive genomic representations.

The current standard genome reference for Homo Sapiens, GRCh38, released in 2013(3), incorporates significant improvements over previous builds, including better assembly quality, updated gene annotations, and corrections to errors found in earlier versions(4). For example, compared to GRCh37, version 7 of GRCh38, named as GRCh38.p7, exhibits modifications in approximately 8000 nucleotides, rectifying misassembled regions, filling in gaps, incorporating additional sequence information for centromeres, and notably enhancing the diversity of the genome reference by integrating 261 alternate loci across 178 distinct regions (Homo sapiens genome assembly GRCh38.p7). GRCh38.p7 has about 358 replacement fragments with a total length of about 109 million bases and spans 60Mb of the primary assembly. These alternate-contigs represent a diversity of human genomes mined from approximately 50 different libraries. Most of GRCh38’s improvements are the result of other genome sequencing and analysis projects, including the 1000 Genomes Project. Beyond fixing thousands of small fragments that led to missing variant calling, GRCh38 specifically adds ALT contigs to describe common complex variations. These contigs represent DNA sequence fragments that differ significantly between different populations. The effectiveness and improvements of GRCh38 led to better accuracy of read alignment and variant calling which have been recognized in many studies(5; 6)

The genome reference is utilized at almost every step of the processing pipeline, however, the current standard genome reference still maintains some limitations. The major part of the human genome reference is genomic sequencing data synthesized from a group of donors from the United States. 65% of the genome reference is built based on the genome of an individual who is 57% European and 37% African(2). Therefore, other populations that are genetically different from Europeans and Africans are not adequately represented in the genome reference. It affects the performance of the statistical analysis between the healthy group and the disease group, as causal pathogenic variants are biased towards genome references built based on different genetic populations from their own.

Many studies in different countries have created entirely new genome references such as Korea(7), Denmark(8), or Egypt(9). It is known that integrating population variants into the standard reference allows one to take advantage of known variant information to improve the sensitivity to detect variants(10). Several studies have built population-specific genome references by this way, such as those for Japan(11) or the United Arab Emirates(12). Both studies built population-specific genome references based on the old standard genome reference GRCh37 provided by the Genome Reference Consortium in 2009. These prior approaches involve substituting common alleles directly within the primary genome (chromosomes 1-22, X, Y, and M) of GRCh37, without addressing other genomic regions(12; 13). This methodology proves advantageous due to the limited presence of alternate sequences in comparison to the primary chromosome in GRCh37. The latest version of GRCh38 has increased to thousands of alternate contigs. These alternate fragments differ from the main chromosome in one part throughout their entire length and remain largely unchanged. However, to the best of our knowledge, no studies consider building a population-specific genome reference that efficiently integrates variants from large-scale population genomics databases into GRCh38 with an awareness of the alternate contigs to take their full advantages.

In this study, we proposed a framework to build a population-specific genome reference using GRCh38 as backbone with alternate-contig awareness to increase variant calling accuracy and demonstrated its usage with Vietnamese genomic data. In particular, we built a genome reference called VGR (**V**iet**n**amese-specific **G**enome **R**eference) by integrating variant information derived from high-coverage whole-genome sequencing (WGS) data of the 1000 Vietnamese Genomes Project (VN1K)(14) into GRCh38 with alternate contigs. VGR was then compared to GRCh38 in genotype calling analyses using additional high-coverage WGS data of 99 Vietnamese individuals (KHV) from the 1000 Genomes Project (1KGP)(15). Our results showed that VGR significantly reduced singleton and supplementary mappings and decreased the complexity of detecting variants compared to GRCh38. Using Omni array genotyping data of 99 KHV samples as a benchmark, we also observed lower false-positive and false-negative rates of variant calling with VGR compared to GRCh38. Given its advantage, this framework could easily be adapted to other populations as long as there is a variant database for them.

## 2. Methods

### 2.1. Framework overview

The proposed framework for constructing a population-specific genome reference utilizing GRCh38 as a backbone is shown in **Figure 1**. The outcome is a population-specific genome reference that represents the population characteristics by integrating common alleles (frequencies greater than 50%) of the target population into the standard genome reference. This approach retained the valuable replacement fragments provided in GRCh38 to reduce errors in variant calling. This method involves breaking down each chromosome of GRCh38 into a set of different fragments and replacement/insertion/deletion of variants can be performed on these component fragments. Using this, the modification will be calculated with a smaller base unit. Other alternate regions of primary contigs are also modified on identical sequence fragments. We calculate the new positions of alternate contigs by changing the positions of the overlapped fragments with alternate fragments. The forward/reverse direction of the compound fragments is used to determine the base to be adjusted.

**Figure 1.**
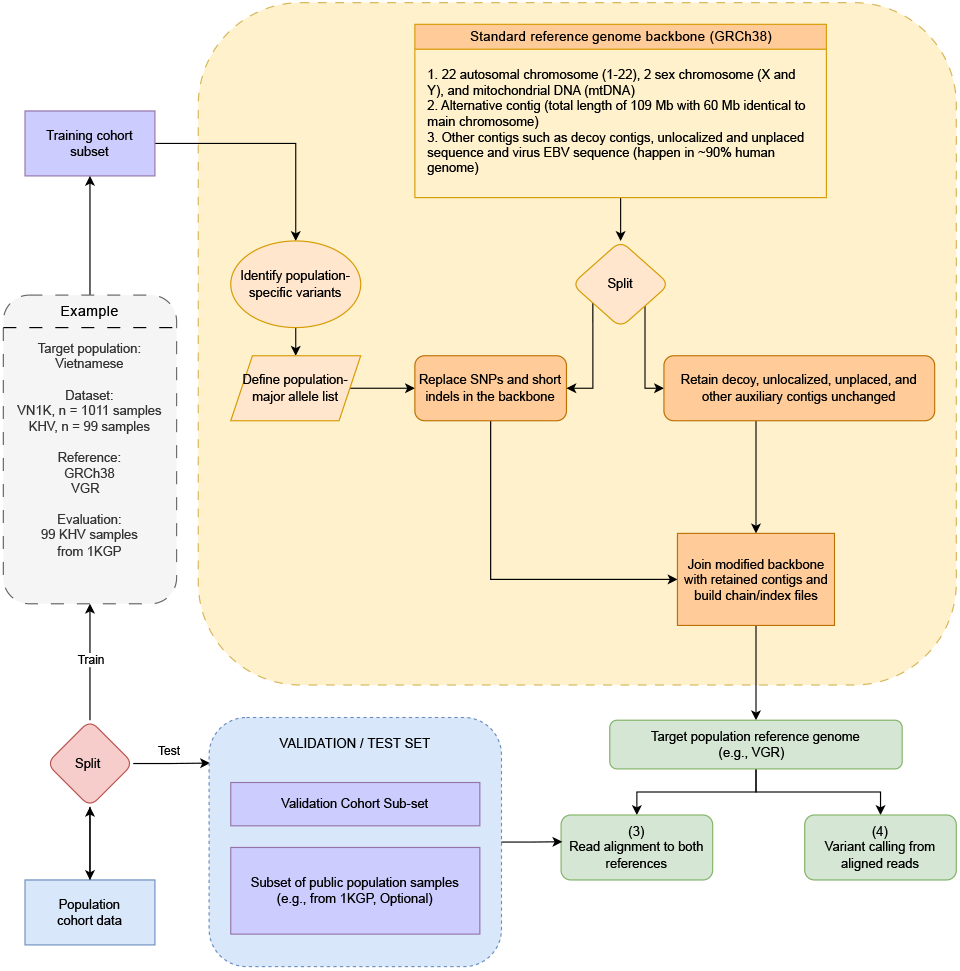
General workflow for building a population-specific genome reference utilizing GRCh38 as a backbone with alternate contigs. The Vietnamese data used in this study is shown as an example.

**Figure 2.**
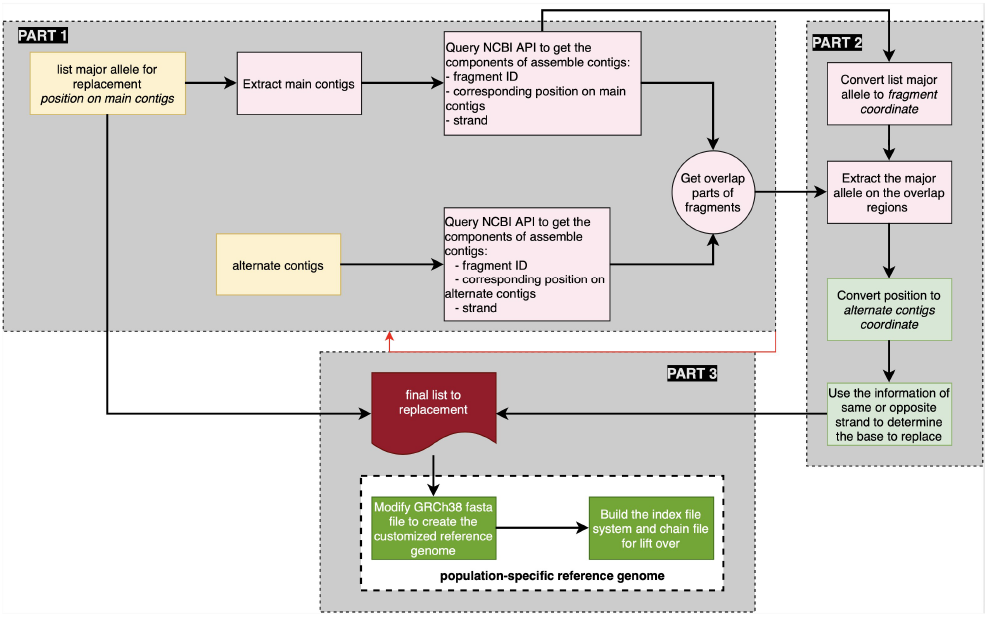
Workflow for handling alternate contigs in building a population-specific genome reference.

### 2.2. Handling alternate contigs in genome reference construction

GRCh38 contains numerous alternate contigs representing structurally divergent haplotypes of genomic regions in the primary assembly. The pipeline for customizing the genome reference consists of three major parts, as outlined in **Figure** In **Part 1**, the primary chromosome and alternate contigs of GRCh38 are decomposed into assembly fragments, from which shared and overlapping regions are identified. Each overlapping fragment is defined based on its genomic position and strand orientation (forward or reverse). The input for this step is a list of common population alleles located on the primary chromosomes, including chromosomes 1–22, X, Y, and M (**Algorithm 1**). After extracting the primary regions from the genome reference, we queried the NCBI assembly API to obtain the corresponding assembly fragments. Allele positions on the primary chromosomes were then converted into coordinates relative to the assembled fragments. The same procedure was applied to alternate contigs to generate a corresponding list of alternate fragments. The intersections between these fragment sets represent regions that overlap between the primary chromosome and alternate contigs (**Algorithm 2**).

In **Part 2**, all alleles located within the overlapping regions were remapped onto the corresponding alternate contigs. The overlapping regions identified in **Part 1** serve as the basis for this coordinate conversion process. This step includes three major operations: remapping variants on the primary chromosomes, projecting variants across overlapping regions, and converting coordinates onto alternate contigs. Strand orientation was additionally considered to ensure correct base substitution in reverse-complement regions. The newly calculated coordinates are merged with the original allele list to generate the final set of variants used for the customization of GRCh38 (**Algorithm 3**).

Finally, in **Part 3**, replacement, insertion, and deletion operations were applied to the GRCh38 FASTA reference sequence using the variant list generated in part 2, resulting in the construction of a population-specific genome reference. Other genomic regions, including EBV sequences, decoy sequences, and unknown sequence fragments, remain unchanged. We then generated the indexing system required for downstream analysis and constructed chain files to enable bidirectional coordinate conversion between the customized genome reference and GRCh38.

#### Algorithm 1 calMajorAllele AltRegion

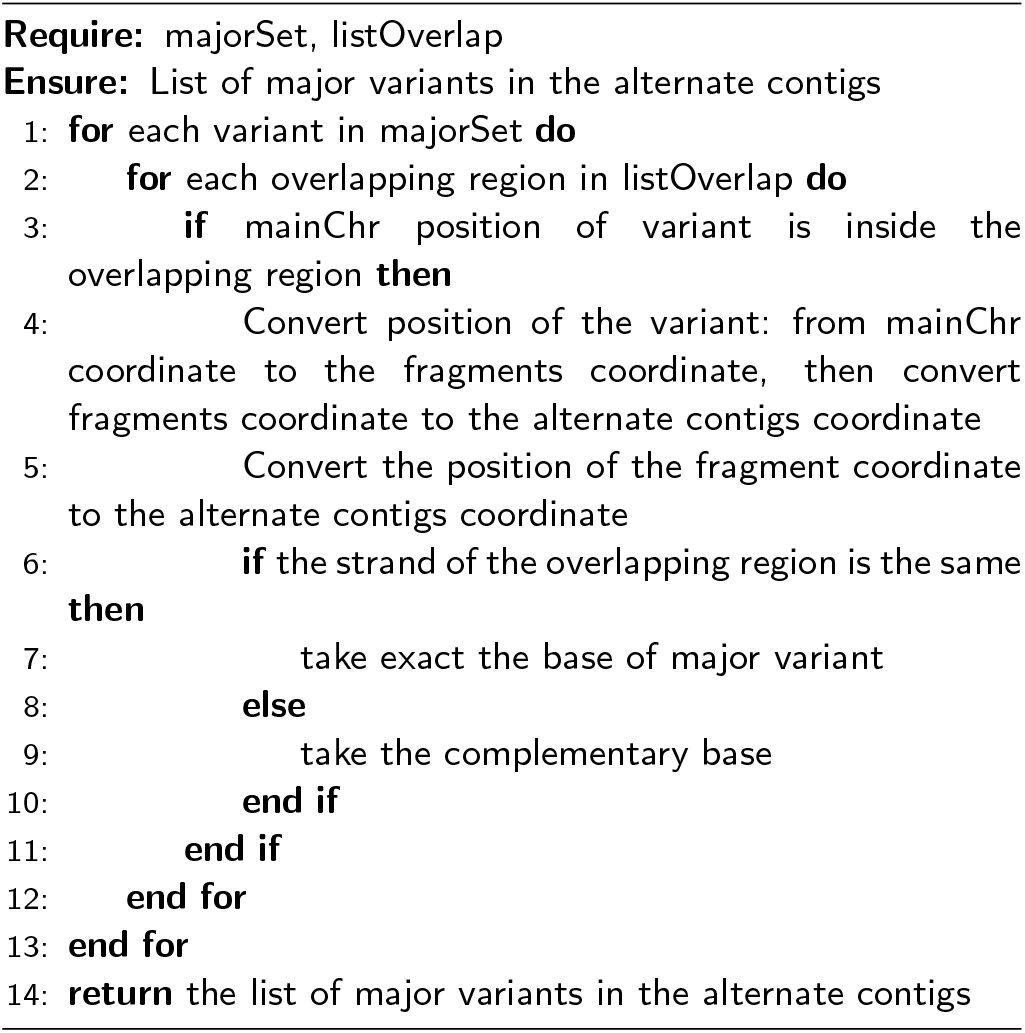

#### Algorithm 2 getChrInfo

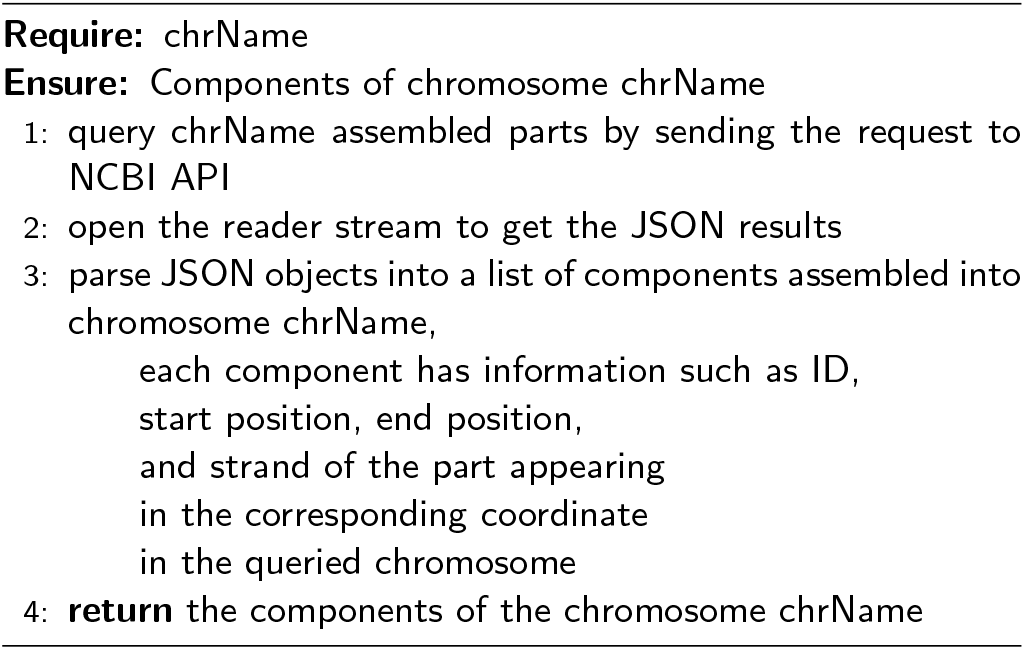

#### Algorithm 3 calOverlapAlterRegion

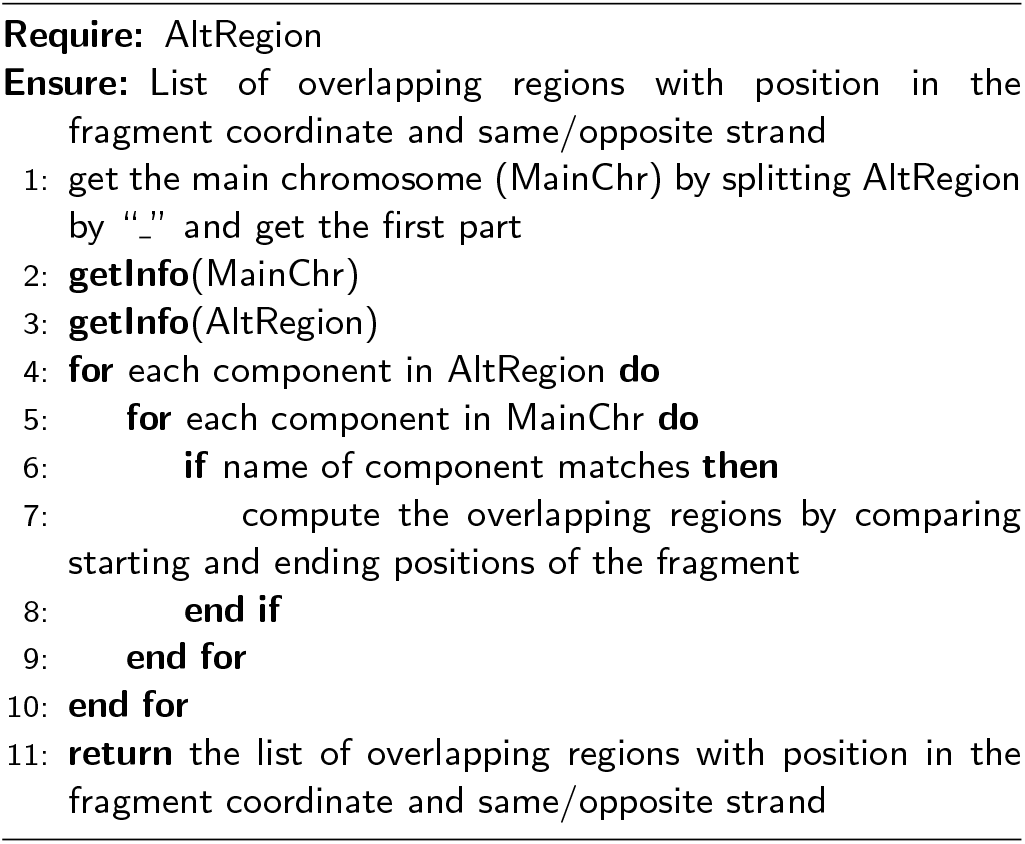

### 2.3. Using the population-specific genome reference

The population-specific genome reference is then integrated into a comprehensive NGS pipeline to enhance alignment and variant calling accuracy, which consists of five primary steps to identify genetic variants: (1) library preparation, (2) sequencing on the Illumina platform, (3) alignment of sequencing reads onto the genome reference, (4) identification of gene variants from these results with or without use of known variation databases, (5) annotation of genomic variants and their function based on different databases (**Figure 3**). In this pipeline, the genome reference serves as the fundamental coordinate system for both read alignment and variant analysis, which is intergrated into step (3) and (4). Finally, to take advantage of the existing genome reference, after step 4 we perform a coordinate conversion (LiftOver) of the identified variants back to GRCh38 coordinate before adding the function annotations in step (5).

**Figure 3.**
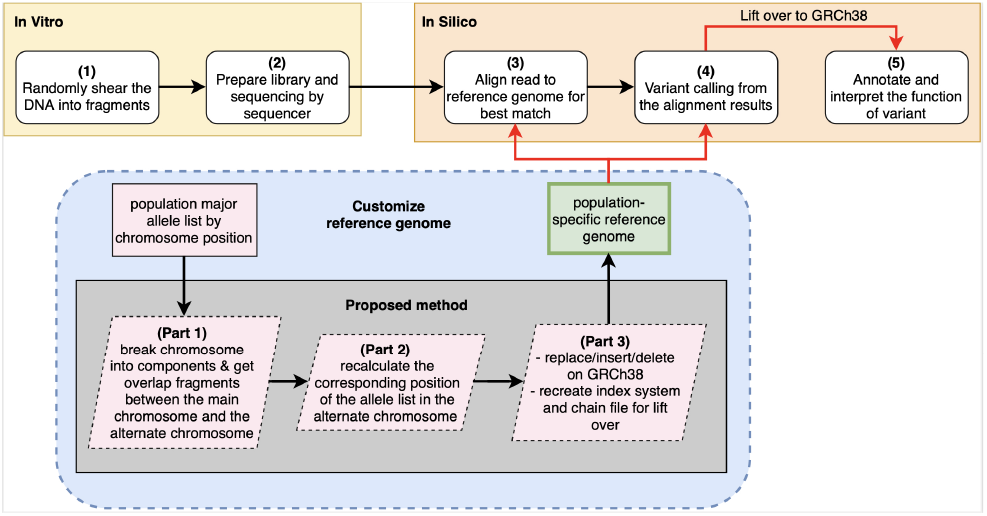
Workflow for data analysis using the population-specific genome reference.

## 3. Results

### 3.1. Experimental setup

The Vietnamese-specific genome reference, VGR, is constructed based on GRCh38 with alternate contigs using VN1K variant information as described in Methods section. To evaluate the performance of VGR in downstream analysis, we used variants derived from high-coverage whole-genome sequencing data of 99 KHV individuals from the 1KGP as an independent test-set. Performance of read alignment and variant calling using VGR was measured by comparing the output to that of GRCh38. **Figure 4** describes this assessment workflow.

**Figure 4.**
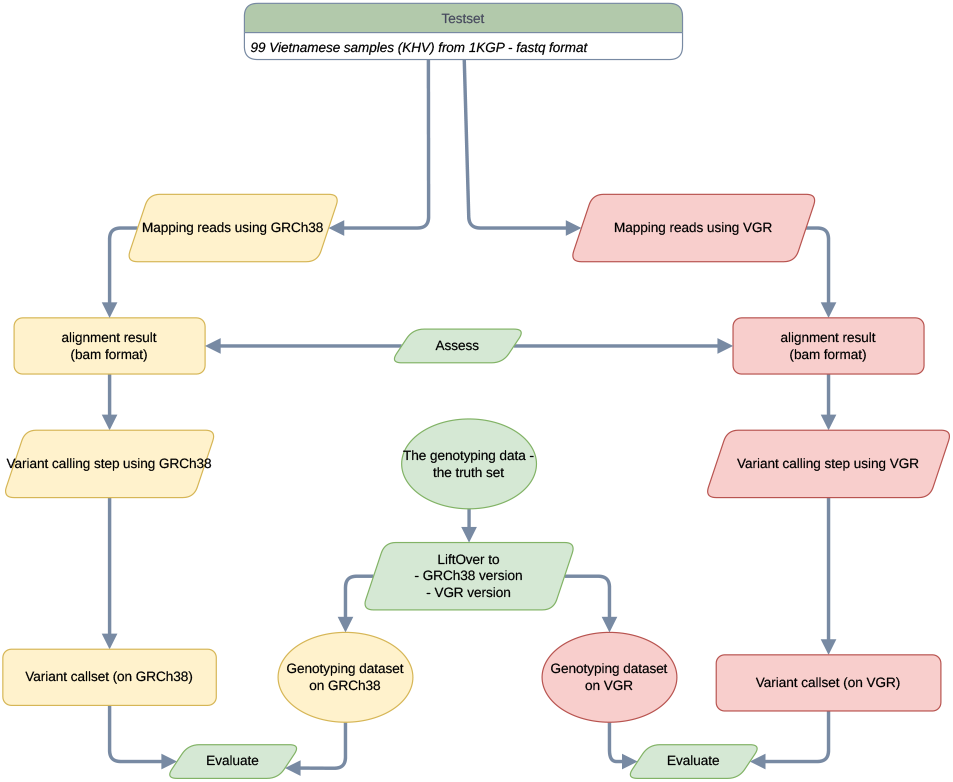
Assessment workflow for VGR and GRCh38 using KHV whole-genome sequencing datasets.

For assessment of read alignment and variant calling, high-coverage WGS reads from the 99 KHV samples from 1KGP were aligned separately to GRCh38 and VGR using NVIDIA Clara Parabricks pbrun germline version 2.4.5. The framework workflow performs read alignment, coordinate sorting, and variant calling in a single pipeline. Assessment of the alignment output (BAM files) was performed using Picard CollectAlignmentSummaryMetrics. For each sample and reference, mismatch rate and high-quality error rate were extracted from the alignment summary metrics. These metrics were compared between GRCh38 and VGR across the same 99 KHV samples using paired statistical tests. Quality of the variant calls (VCF files) was then assessed using Picard CollectVariantCallingMetrics. Additional VCF processing, filtering, and summary analyses were performed using bcftools and samtools. The variants called against VGR were converted back to GRCh38 coordinates using GATK LiftoverVcf with the VGR-to-GRCh38 chain file generated during the reference construction. For population structure analysis, filtered autosomal biallelic SNPs were converted to PLINK format, linkage-disequilibrium pruning was performed, and principal component analysis was conducted using PLINK.

The Omni array genotyping data from the 1KGP were used as an independent benchmark to evaluate precision and recall of variant calls from VGR and GRCh38. For each KHV sample, WGS-derived variant calls generated using GRCh38 and VGR were compared with Omni genotype calls at shared variants. True positives (TP) were defined as concordant variant genotypes between the WGS callset and Omni data, false positives (FP) as WGS variant calls not supported by Omni genotypes, and false negatives (FN) as Omni variant genotypes missed by the WGS callset. True negatives (TN) were not considered because the comparison focused on variant sites represented in the Omni benchmark rather than all homozygous-reference positions across the genome.

### 3.2. Integrated variant landscape of VGR

To characterize the sequence-level modifications incorporated in VGR, we compared it directly with the GRCh38 backbone using the chromosome-wise reference-to-reference alignment shown in **Figure 5**. This analysis identified 1,910,933 reference differences across primary chromosomes, including 1,868,232 SNPs, 24,983 insertions, and 17,718 deletions. SNPs accounted for 97.8% of all inferred differences. Genome-wide density analysis further showed that these differences were broadly distributed across chromosomes, with several local peaks in 1-Mb windows.

**Figure 5.**
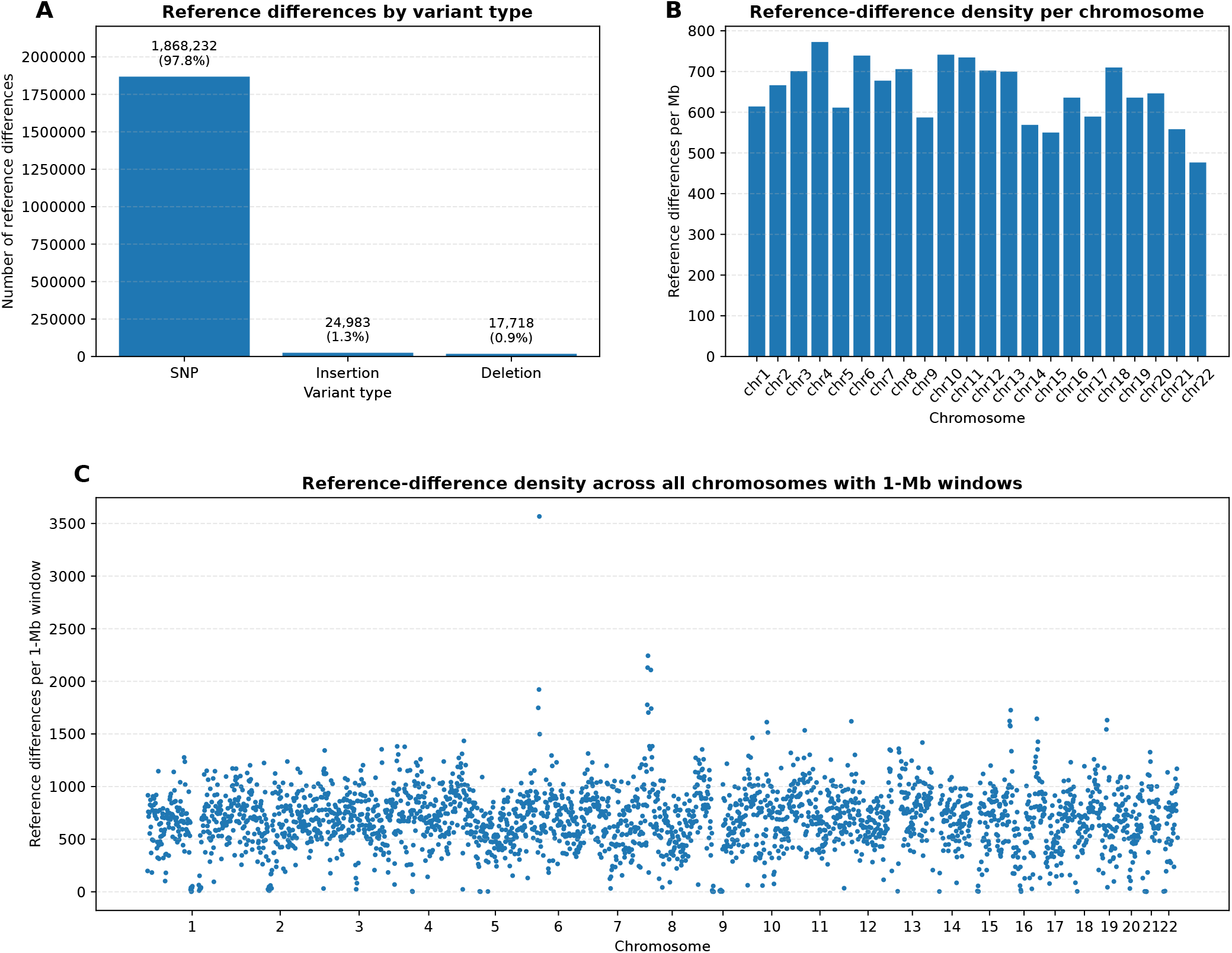
Sequence differences between the final VGR and the GRCh38 were identified using chromosome-wise FASTA-to-FASTA alignment. (A) Number of reference differences between VGR and GRCh38 by variant type, including SNPs, insertions, and deletions. (B) Chromosome-level density of reference differences between VGR and GRCh38, normalized by chromosome length and shown as reference differences per megabase. (C) Genome-wide distribution of reference differences between VGR and GRCh38 across 1-Mb windows

### 3.3. VGR improves read alignment

Read alignment quality was evaluated using mismatch rate and high-quality (HQ) error rate metrics across the 99 KHV samples. VGR demonstrated significantly lower mismatch and HQ error rates compared with GRCh38 (Wilcoxon signed-rank test, p ¡ 0.001 for both metrics) (**Figure 6**). Specifically, the median mismatch rate decreased from approximately 0.0040 with GRCh38 to 0.0037 with VGR, while the median HQ error rate decreased from approximately 0.0033 to 0.0029. These improvements were consistently observed across most evaluated samples, indicating that VGR provided more stable alignment performance than GRCh38.

**Figure 6.**
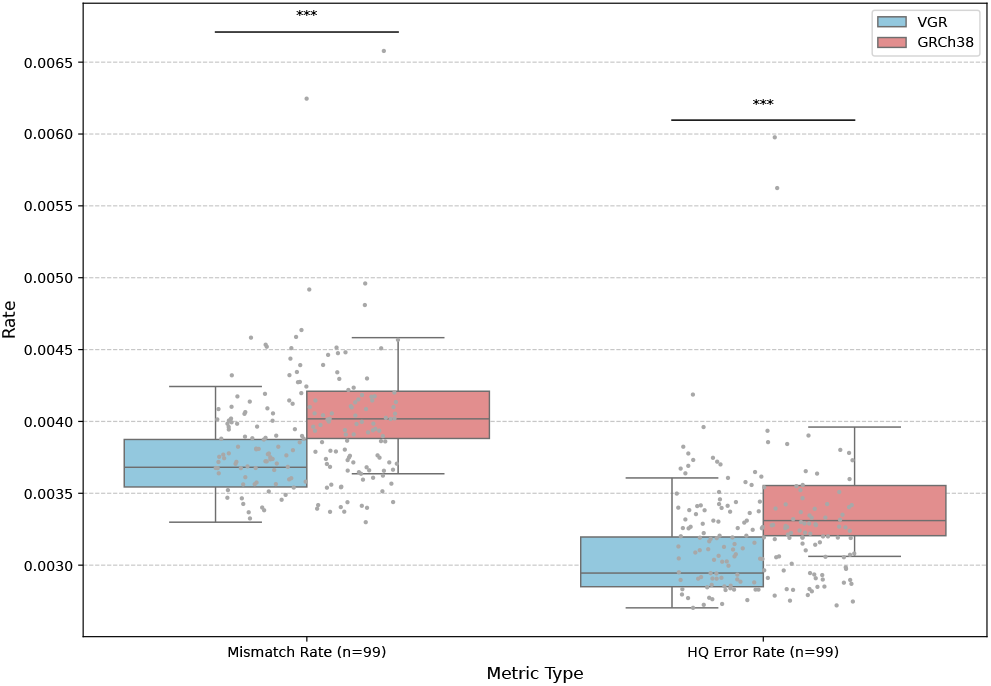
Comparison of mismatch and high-quality (HQ) error rates of variant calls using GRCh38 and VGR across 99 KHV whole-genome sequencing datasets. Statistical significance was assessed using the Wilcoxon signed-rank test (p-value*<*0.05: *; *<* 0.01: **; *<* 0.001: ***).

The improved alignment quality observed with VGR likely reflects the incorporation of Vietnamese variants into the standard genome reference. By reducing the differences between sequencing reads and the reference sequence, VGR decreased alignment mismatches and reduced ambiguity during read mapping. This effect is particularly important in regions containing population-specific variants, where the standard genome reference may introduce reference bias and increase mapping errors. The reduction in HQ error rates further suggests that reads aligned to VGR were supported by higher-confidence base calls, potentially improving the reliability of downstream variant detection analyses.

### 3.4. Variant calling evaluation

Variant calling performance was evaluated for VGR and GRCh38 using Omni array genotyping data as an independent genotyping benchmark (**Figure 7**). VGR achieved substantially higher precision together with fewer FP calls, while GRCh38 produced a slightly higher absolute number of TP calls. Specifically, the median precision increased from approximately 0.972 with GRCh38 to 0.995 with VGR, while the median FP count decreased from approximately 1200 to 1100. These findings indicate that VGR reduced erroneous variant calls and improved the specificity of variant detection relative to GRCh38. The reduced number of TP calls observed in VGR likely reflects the lower overall number and complexity of the detected variants after incorporation of population-specific alleles into the customized genome reference. Because many common Vietnamese alleles were directly integrated into VGR, a subset of previously identified variants relative to GRCh38 became reference-matching positions in the customized genome. Consequently, the total number of detected variants was reduced, leading to fewer overall TP calls while simultaneously reducing FP calls and improving precision metrics.

**Figure 7.**
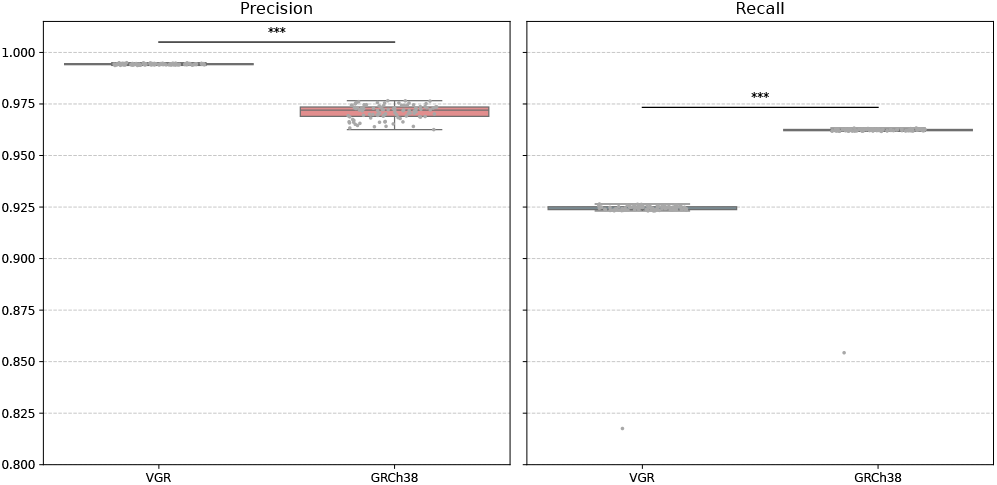
Comparison of precision and recall of variant calls using VGR and GRCh38 across 99 KHV whole-genome sequencing datasets using Omni array genotyping data as independent genotyping benchmark. Statistical significance was assessed using the Wilcoxon signed-rank test (p-value¡0.05: *; ¡0.01: **; ¡0.001: ***).

In contrast, GRCh38 achieved higher recall values, with a median recall of approximately 0.962 compared to 0.925 for VGR. Although VGR showed a single FN outlier exceeding 60,000, the overall FN distributions remained relatively similar between the two groups. These findings suggest that VGR adopts a more conservative variant-calling behavior, prioritizing the reduction of ambiguous and potentially erroneous variant calls while maintaining comparable sensitivity across most samples. Moreover, We compared the minor allele frequency distributions of filtered biallelic SNPs from GRCh38 and VGR callsets in the 99 KHV samples (**Figure 8**). Because the total number of retained variants differed between references, MAF distributions were plotted as the percentage of SNPs within each callset. The GRCh38 callset showed a higher proportion of low-frequency variants, particularly near MAF = 0%. In the rare-variant range, both callsets showed a peak around 0.5%, corresponding to singleton alleles among 99 diploid samples. These patterns suggest that incorporation of Vietnamese-major alleles into VGR changes the frequency spectrum of retained variant calls, reducing the relative contribution of low-frequency calls and enriching for common polymorphisms among variants still detected relative to the customized reference.

**Figure 8.**
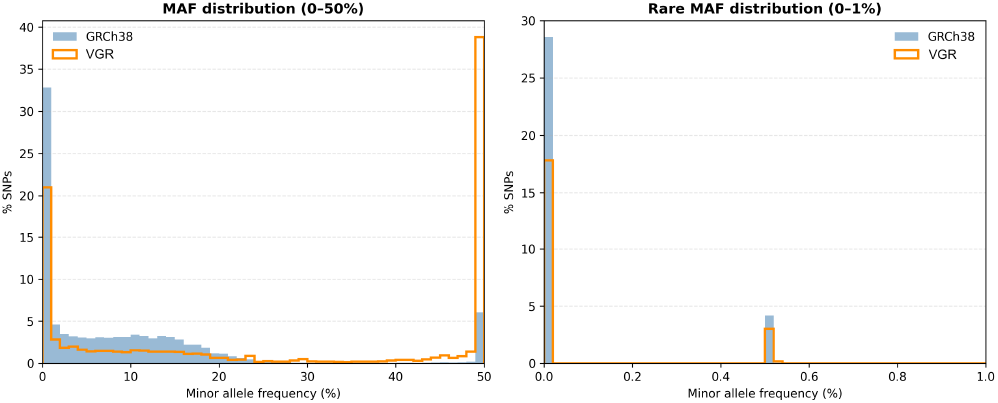
MAF distribution of 99 KHV samples calculated with GRCh38 and VGR

To further evaluate whether the use of VGR affected population-level analyses, we performed principal component analysis (PCA) on Asia populations. The PCA clustering patterns were highly concordant between the callsets aligned with VGR and GRCh38 (**Figure 9**). In both analyses, KHV samples appeared near CDX samples in both PCA plots. JPT samples formed a separate group, while CHB and CHS samples partly overlapped. The very high correlations between the two results of PCA (PC1: r = 0.99999; PC2: r = 0.99993) show that the use of VGR did not noticeably change the population structure compared to GRCh38.

**Figure 9.**
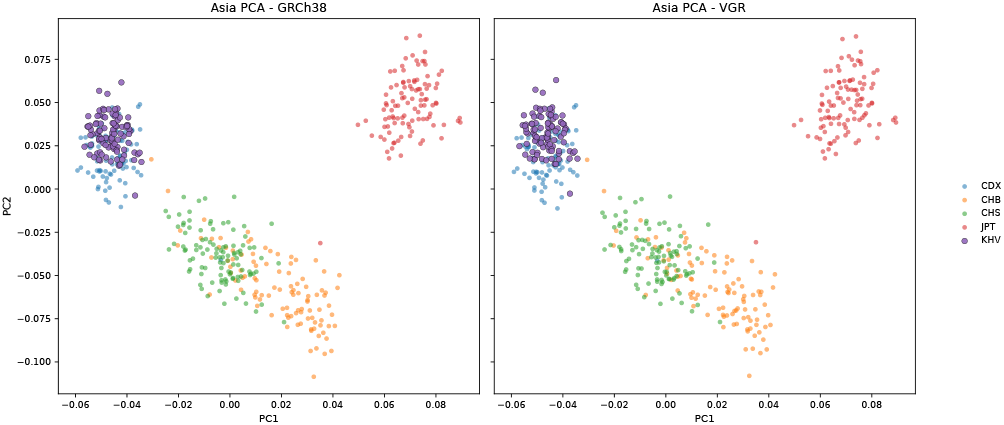
Principal component analysis of Asian populations using GRCh38- and VGR-derived variant callsets. PCA was performed using autosomal biallelic SNPs followed by LD pruning. The two references produced highly concordant clustering patterns, indicating that VGR preserved population-level genetic structure relative to GRCh38.

## 4. Discussion

In this study, we proposed a framework for constructing a population-specific genome reference based on GRCh38 as a backbone with explicit handling of alternate contigs. By integrating variants from the Vietnamese population using the VN1K dataset into the genome reference, we generated VGR and evaluated its performance using WGS and genotyping data of 99 KHV samples from the 1KGP. Our results provided a Vietnamese-specific linear reference that reduced reference mismatch during read alignment, improved variant-calling precision, altered the spectrum of retained variant calls, while preserving population-level genetic structure.

This study has several limitations. First, the evaluation was performed primarily using KHV whole-genome sequencing datasets from the 1000 Genomes Project, which may not fully represent the genetic diversity of all Vietnamese subpopulations. Second, the benchmarking was based on Omni genotyping array datasets, which evaluated only a subset of genomic variants and may not fully capture structural variants or rare variants. Third, although VGR improved precision and reduced FP calls, a moderate reduction in recall was also observed, indicating that some true variants may become more difficult to detect using a highly population-specific genome reference.

Future work should expand the construction and evaluation of Vietnamese population-specific genome references using larger and more diverse cohorts and data, including long-read sequencing data and graph-based genome representations. Integration of structural variants and haplotype-aware genomic models may further improve the representation of complex genomic regions beyond the capabilities of linear genome references. In addition, a VGR evaluation will be performed in clinical and disease-association studies to help clarify the practical benefits of population-specific genome references for precision medicine applications in Vietnamese populations.

## 5. Data availability

VN1K high-coverage (30×) whole-genome sequencing data was obtained from the MASH Data Portal (genome.vinbigdata.org). Individual-level VN1K data are under managed access and can be requested through the portal in accordance with VN1K Project data access policies. High-coverage (30×) whole-genome sequencing data of KHV samples from the 1000 Genomes Project (1kGP) are publicly accessible through the International Genome Sample Resource (IGSR) data portal (www.internationalgenome.org/data).

## 7. Author contributions statement

N.S.V conceived and supervised the project. T.T.H.T and N.N.N implemented the code. V.C.D and T.H.H participated in result analysis. T.M.P and Q.T.V prepared the datasets. M.H.T and N.T.D performed sample collection and data curation. T.T.H.T and V.C.D drafted the manuscript. N.S.V and Q.N revised analysis results and manuscript. All authors read and approved the manuscript.

## 8. Acknowledgments

We thank internship students at Vingroup Big Data Institute, Hanoi, Vietnam (now reorganized as VinUni Big Data Research Institute, VinUniversity, Hanoi, Vietnam) for their help and feedback in this work.

